# Endosomal GPR65 signaling in fibroblast-like synoviocytes promotes inflammatory cytokine release and nociceptive neuron sensitization

**DOI:** 10.64898/2026.06.16.732753

**Authors:** Luke A. Pattison, Maya Dannawi, Ewan St. John Smith

## Abstract

GPR65 is a proton-sensing G protein-coupled receptor implicated in inflammatory pain. In fibroblast-like synoviocytes (FLS), GPR65 activation promotes the release of proinflammatory cytokines capable of sensitizing sensory neurons. Following stimulation by protons, the synthetic agonist BTB09089, and the glycosphingolipid psychosine GPR65 undergoes internalization; however, the contribution of this trafficking to downstream signaling remains unclear. Using heterologous cell systems, the molecular mechanisms governing GPR65 internalization were first defined. Pharmacological and genetic inhibition of internalization revealed that intracellular trafficking is required for activation of extracellular-signal-related kinase (ERK) in the nucleus and transcriptional responses, indicating a spatially restricted signaling program originating from endosomes. The physiological relevance of this pathway was then examined in primary mouse FLS. Inhibition of endogenous GPR65 internalization reduced the ability of the conditioned media from BTB09089 stimulated FLS to sensitize dorsal root ganglia sensory neurons, thus linking receptor trafficking to pro-nociceptive function. Together these findings identify receptor internalization as a key determinant of nuclear ERK signaling and transcription downstream of GPR65 and demonstrate that endosomal signaling is required for pro-nociceptive activity of GPR65 in FLS.

**One-sentence summary:** Endosomal internalization of GPR65 is required to coordinate gene transcription and proinflammatory cytokine production that drive neuronal sensitization.

## Introduction

The G protein-coupled receptor (GPCR) superfamily is the largest class of receptors targeted by drugs currently approved for use by the US Federal Drug Administration (*1*), however pharmacological understanding of GPCRs is uneven. A particularly neglected GPCR subfamily is the proton-sensing G protein-coupled receptors (PS-GPCRs): six GPCRs tuned to be activated by changes in extracellular pH, with many also activated by various lipid species (*2*). The PS-GPCRs exhibit widespread expression throughout the body and have been linked to several physiological processes, with their dysregulation also contributing to a number of diseases (*3*). However, pharmacological tools to modulate PS-GPCR activity are limited and the family thus represent an untapped resource for development of new therapeutics. Among the PS-GPCRs, GPR65, also known as the T Cell Death Associated Gene 8 (TDAG8) receptor has genetic associations with inflammatory conditions including, ankylosing spondylitis (*4*), chronic obstructive pulmonary disorder (*5*) and inflammatory bowel disease (*6*). Preclinical work directly links expression of GPR65 to worsened pathology in experimental colitis (*7*), and expression and/or activation of GPR65 is concurrent with increased pain-like behaviors in the carrageenan model of inflammatory pain, following intraplantar application of acid (*8*), and in a more translationally relevant preclinical model of rheumatoid arthritis (*9*, *10*). Our own work has described a role of GPR65 on fibroblast-like synoviocytes (FLS) – the cells which line synovial joints – in driving inflammatory responses in the knee joint that cause pain in arthritis (*11*). In addition to protons (*12*), GPR65 is activated by the glycosphingolipid psychosine (*13*) and the synthetic agonist BTB09089 (BTB) (*14*). Our previous systematic characterization of GPR65 agonists revealed that they all evoke receptor internalization, a phenomenon independently observed by others (*13*, *15*). This is particularly noteworthy given the growing interest in the spatiotemporal regulation of GPCR signaling (*16*).

The classical view of GPCR-driven signaling being borne from the plasma membrane, has been eclipsed with growing reports of cells dynamically shuttling GPCRs between the plasma membrane and membranous subcellular compartments (*16*). Often a response to receptor activation, it has been shown that GPCRs not only continue to signal following internalization, but that they drive prolonged signaling events from endosomal platforms (*17*, *18*). Such reports highlight the dynamicity of GPCRs and suggest a possibility of drug development strategies that might enable the decoupling of distinct temporal components of signaling. The sustained signaling of GPCRs from endosomal compartments is of considerable interest within the context of nociception, where it has been shown to drive persistent pain via a number of pronociceptive receptors including protease-activated receptor 2 (*19*), the neurokinin 1 receptor (*20*) and the calcitonin receptor-like receptor (*21*). Further evidence to support the importance of sustained signaling from endosomes in the context of pain, comes from reports that endosomal signaling of the delta opioid receptor confers sustained antinociception (*22*).

The emerging importance of subcellular GPCR signaling in nociception, together with recent reports implicating GPR65 in inflammatory pain and its agonist-driven internalization, highlights an important unresolved question: how does the spatial regulation of GPR65 shape its signaling output? Here, the molecular mechanisms governing GPR65 internalization are defined, disrupting receptor internalization in a stable cell line revealed compartmentalized signaling events driven by endosomal GPR65. Extending these findings to endogenous systems, the subcellular location of GPR65 was shown to dictate downstream signaling in FLS, affecting physiologically relevant outputs including transcriptional programs and paracrine communication, which ultimately drive sensitization of nociceptive neurons. Collectively, these findings identify spatial encoding as a key mechanism underlying GPR65 function in pain and highlight endosomal signaling of GPCRs in FLS as an important determinant in nociceptive pathways.

## Results

### Upon activation GPR65 internalizes to early endosomes in a dynamin and clathrin dependent manner

GPR65 has previously been shown to leave the plasma membrane upon activation by protons, BTB and psychosine (*11*, *13*, *15*). Agonist-induced internalization was further confirmed in Flp-In CHO cells using confocal microscopy to assess the localization of mouse GPR65 with a C-terminal mCherry tag (mGPR65-mCherry) following 30-min stimulation with GPR65 agonists or vehicle controls. For cells exposed to neutral Hanks Balanced Salt Solution (HBSS; pH 7.4) or 0.1% (v/v) DMSO, mCherry fluorescence was predominantly observed at the cell surface boundary, whereas exposure to acidic HBSS, pH 6.0) or 100 µM BTB resulted in the emergence of intracellular fluorescent puncta, indicative of receptor internalization (Fig. 1A). The number of intracellular puncta was shown to be dependent on treatment (*H*(4,110) = 61.161, *p* < 0.0001), with significant increases for test solutions over their corresponding vehicle (HBSS pH 7.4, 0.021 ± 006, HBSS pH 6.0, 0.111 ± 0.018, Z = 4.04, *p-adj =* 0.0008; DMSO, 0.015 ± 0.002, BTB, 0.104 ± 0.017, Z = 5.09, *p-adj* < 0.0001; Fig. 1B). To better identify the subcellular location of mGPR65 following internalization, bioluminescence resonance energy transfer (BRET) was assessed between luciferase tagged GPR65 (mGPR65-RLuc8) and Venus tagged Rab5a (Rab5a-Venus), a marker of early endosomes (Fig. 1C). Consistent with previously reported decreases in BRET between mGPR65-RLuc8 and a plasma membrane marker (*11*), an increase in BRET signal between mGPR65-RLuc8 and Rab5a-Venus was measured upon stimulation with acidic buffer solutions (Fig. 1D), or BTB (Fig. 1E) indicative of internalization to early endosomes. The agonist-induced internalization of mGPR65 to early endosomes was shown to be concentration-dependent, with EC_50_ estimates of pH 6.77 and 11.4 µM for protons and BTB respectively (Fig. 1F). Similar results were obtained when studying the influence of psychosine-stimulation of cells expressing mGPR65-mCherry or the mGPR65-RLuc8, Rab5a-Venus BRET pairing (Fig. S1).

**Figure 1.**
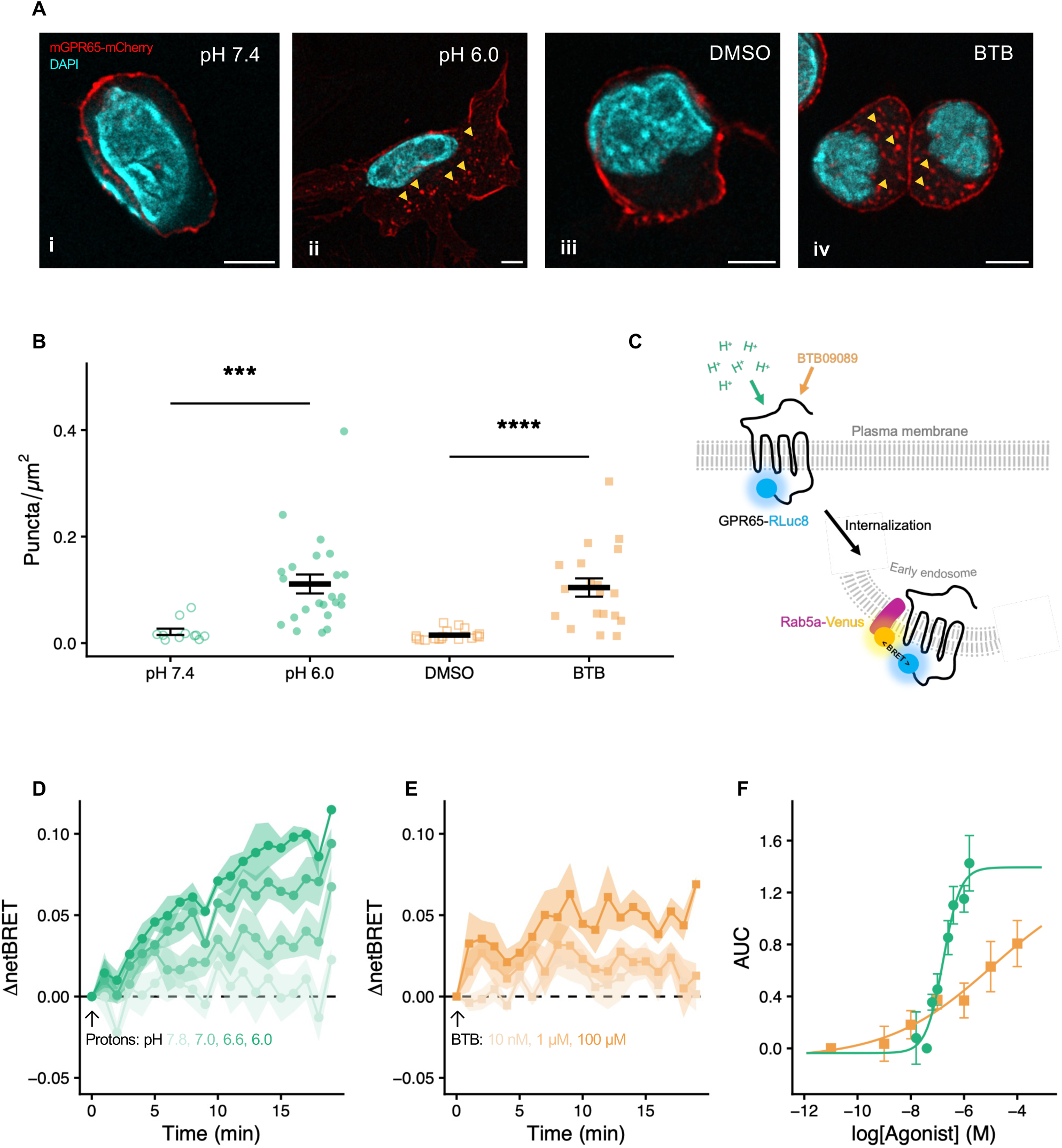
Following activation GPR65 internalizes to early endosomes. **(A)** Representative images of Flp-In CHO cells expressing GPR65-mCherry following 30-min stimulation with **(i)** HBSS (pH 7.4), **(ii)** HBSS (pH 6.0), **(iii)** 0.1% (v/v) DMSO or **(iv)** 100 µM BTB (red, GPR65-mCherry; blue: nuclear stain; yellow arrowheads: example intracellular puncta; scale bar, 5 µm). **(B)** Quantification of intracellular fluorescent puncta normalized to total cell area. **(C)** Bioluminescence resonance energy transfer (BRET) between GPR65-RLuc8 and Rab5a-Venus was used to quantify receptor localization to early endosomes. **(D, E)** GPR65 internalization following stimulation with **(D)** protons (circles) or **(E)** BTB (squares). **(F)** Concentration-response analysis of agonist-induced internalization. Data in **(B)** represent individual cells from three independent experiments; data in **(D-F)** represent means from four independent experiments. *** *p-adj* < 0.001, **** *p-adj* < 0.0001; Kruskal-Wallis test (stimulation) with Bonferroni-corrected post hoc.

In combination with other proteins, GPCR endocytosis is mediated by dynamin GTPases and clathrin; clathrin driving the formation of endocytic pits (*23*), which are later cleaved from the membrane by dynamin (*24*). Disrupting these proteins via pharmacological and genetic mechanisms has been shown to be an effective strategy to prevent GPCR internalization, validated for several GPCRs in similar BRET systems to that used in Fig. 1A (*19*, *25–27*). To assess the dynamin dependency of mGPR65 internalization to early endosomes, BRET assays were repeated in cells overexpressing a dominant negative mutant of dynamin, Dynamin K44E (DynK44E), which is deficient in GTPase activity (*28*); overexpression of wild-type dynamin (DynWT) and mock transfection (pcDNA3.1) served as controls. Clathrin-dependency was investigated by pre-treating cells with the clathrin inhibitor, PitStop2 (PS2; Fig. 2A). For both strategies a significant effect of experimental condition was observed (Dynamins: *F*(2,39) = 61.789, *p* < 0.0001; PS2: *F*(1,24) = 121.517, *p* < 0.0001). Overexpression of DynK44E significantly attenuated the ability of mGPR65 to associate with Rab5a-Venus expressing early endosomes in response to proton-stimulation, with this interaction not affected in mock transfected cells or cells expressing DynWT (AUCs: pcDNA3.1, 1.178 ± 0.189, DynWT, 1.035 ± 0.134, DynK44E, −0.315 ± 0.161, *F*(2,39) = 47.5, *p-adj* < 0.0001; Fig. 2B). Similar results were obtained when stimulating cells with 100 µM BTB (pcDNA3.1, 0.485 ± 0.067, DynWT, 0.605 ± 0.060, DynK44E, −0.190 ± 0.090, *F*(2,39) = 10.7, *p-adj = 0.0002*; Fig. 2C). The abilities of protons and BTB to trigger mGPR65 internalization to early endosomes was also negatively affected by PS2 inhibition of clathrin, but not DMSO vehicle control (Protons: DMSO, 1.480 ± 0.144, PS2, 0.574 ± 0.093, *p-adj* = 0.0007; Fig. 2D; BTB: DMSO, 0.644 ± 0.078, PS2, −0.173 ± 0.095, *p-adj* = 0.0002; Fig. 2E). Similarly, psychosine-induced internalization of GPR65 was also shown to be dependent on dynamin and clathrin (Fig. S2).

**Figure 2.**
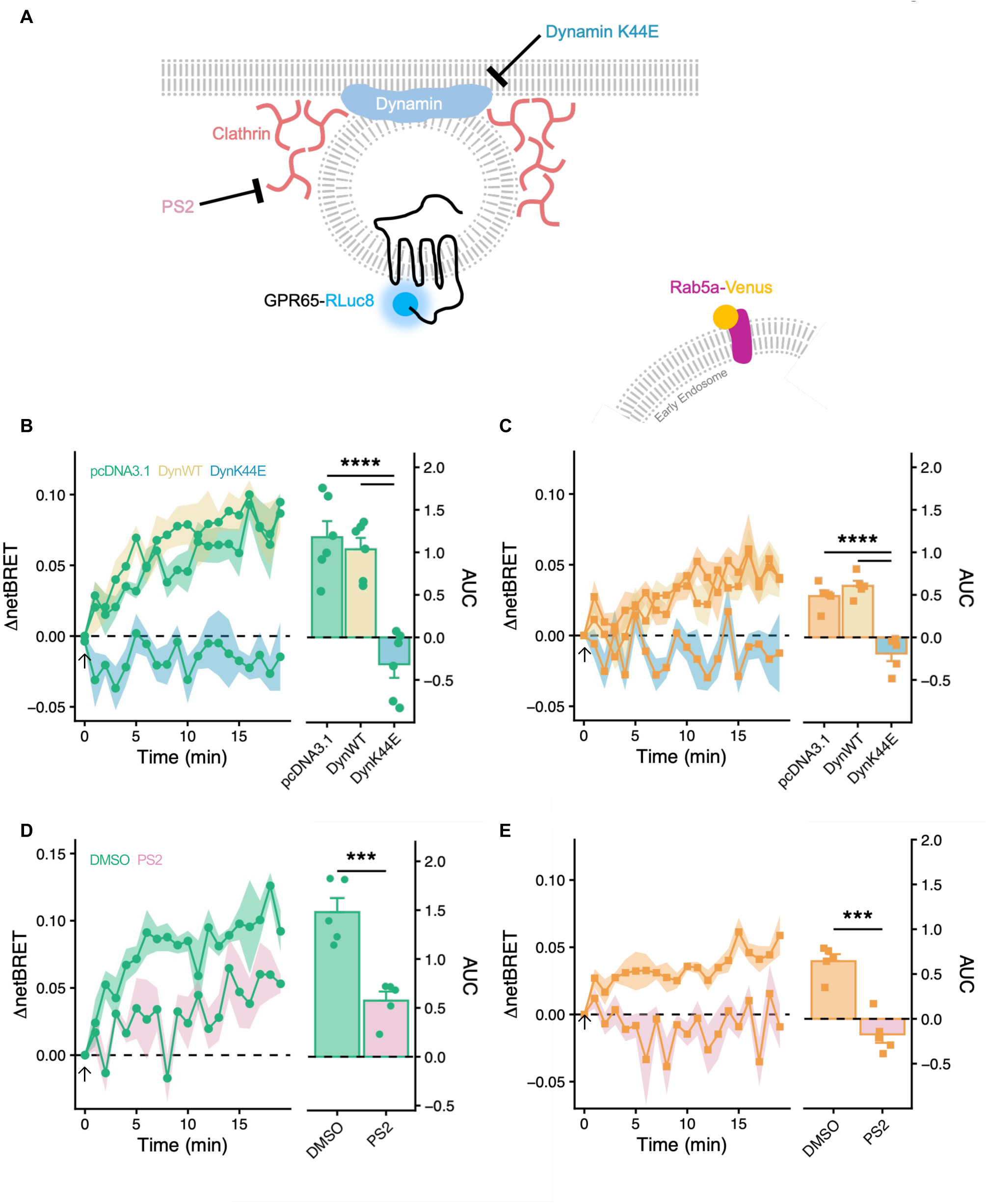
GPR65 internalization is dependent on dynamin and clathrin. **(A)** GPCR internalization is mediated by dynamin GTPases and clathrin and can be inhibited pharmacologically (PitStop2, a clathrin inhibitor; PS2) or genetically by overexpression of a GTPase-deficient dominant-negative dynamin mutant (DynK44E). **(B, C)** BRET between GPR65-RLuc8 and Rab5a-Venus was measured in Flp-In CHO cells overexpressing DynK44E, wild-type dynamin, or pcDNA3.1 upon stimulation with **(B)** HBSS (pH 6.0; circles), or **(C)** 100 µM BTB (squares). **(D, E)** Effect of clathrin inhibition on GPR65 internalization following stimulation with **(D)** HBSS (pH 6.0; circles) or **(E)** 100 µM BTB (squares) in cells pre-treated with 30 µM PS2 or 0.1% (v/v) DMSO. Data represent five independent experiments. *** *p-adj* < 0.001, **** *p-adj* < 0.0001; two-way ANOVA (stimulation x condition) with Bonferroni-corrected post hoc.

### Inhibiting internalization of GPR65 to early endosomes has a knock-on effect on the downstream signaling consequences that can be elicited

The interplay between GPCR subcellular localization and signaling output is of emerging interest, with increasing reports of subcellularly localized GPCRs driving distinct signaling processes, reviewed in (*18*). To investigate the contribution of mGPR65 located at early endosomes to signaling output, endpoint signaling assays were performed on cells overexpressing DynK44E, or mock transfected with pcDNA3.1. GPR65 predominantly couples to Gα_s_ proteins and, accordingly, stimulation of CHO cells stably expressing mGPR65 with pH 6.0 or 100 µM BTB coordinated cAMP accumulation, which was reduced in DynK44E-expressing cells (*F*(1,12) = 28.408, *p* = 0.0002; Fig. 3A). Importantly, DynK44E did not affect the cAMP generated by forskolin treatment, which was used to normalize data (pcDNA3.1, 3,364.0 ± 654.7 RFU, DynK44E, 3,954.3 ± 570.9 RFU, t = 0.679, df = 6, *p* = 0.522; Fig. S3A). Generation of cAMP is necessary for activation of protein kinase A, the longer-term consequences of activating a Gα_s_-coupled receptors being driven by protein kinase A and relating to phosphoregulation of effector proteins, including the cAMP response element (CRE). Thus, CRE activity in response to GPR65 activation was also investigated using a CRE-dependent nanoluciferase (NLuc) reporter system (*29*). Owing to the 6-h stimulation of these experiments only BTB was used to activate GPR65. Parental Flp-In CHO cells were transfected with the CRE-NLuc reporter, mGPR65-mCherry and either DynK44E or pcDNA3.1. The comparable intensity of mCherry fluorescence between cells expressing DynK44E and pcDNA3.1 (pcDNA3.1, 100,793.1 ± 2,342.3 RFU, DynK44E, 100,254.2 ± 2,064.9 RFU, t = −0.173, df = 6, *p* = 0.869; Fig. S3B) indicated similar transfection efficiencies and suggests that the attenuated CRE induction of NLuc in cells expressing DynK44E is most likely due to the impaired ability of GPR65 to internalize (pcDNA3.1, 2.21 ± 0.08, DynK44E, 1.51 ± 0.12, t = −4.741, df = 6, *p* = 0.0032; Fig. 3B). GPR65 activation also stimulates extracellular-signal regulated kinases (ERK): fitting with previous results in cells where GPR65 could not internalize, both protons and BTB were impaired in their activation of ERK (*F*(1,12) = 23.044, *p* = 0.0004; Fig. 3C). The ability of the positive control, PDBu, to activate ERK was not affected by DynK44E expression (pcDNA3.1, 4,718.9 ± 1,136.2 RFU, DynK44E, 5,547.5 ± 1,147.8 RFU, t = 0.513, df = 6, *p* = 0.626; Fig. S3C).

**Figure 3.**
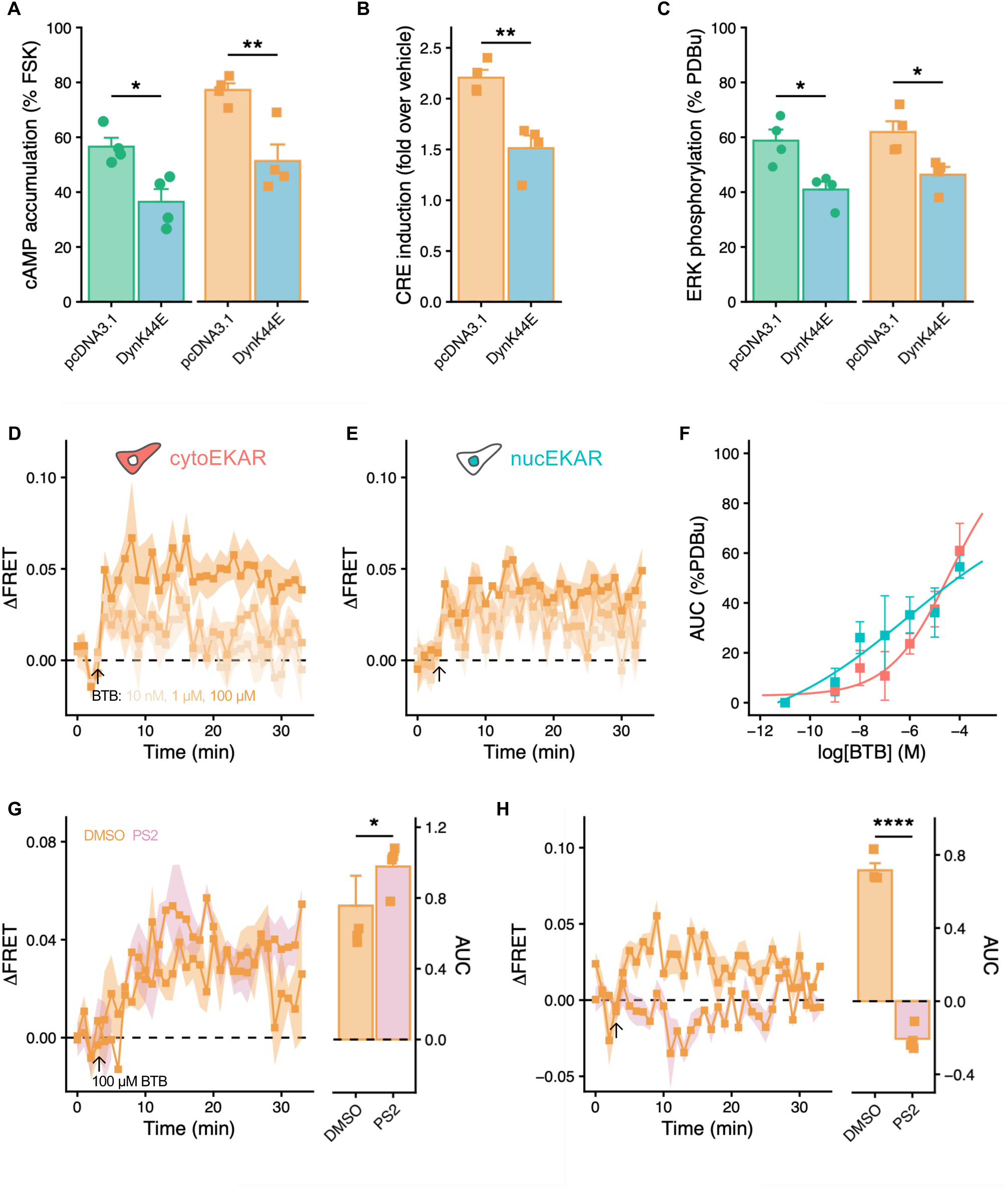
GPR65 internalization shapes signaling output in CHO cells. **(A-C)** The effect of DynK44E overexpression on **(A)** pH 6-(circles) and 100 µM BTB-(squares) induced cAMP production, **(B)** 100 µM BTB-induced CRE-NLuc reporter activity, and **(C)** pH 6-(circles) and 100 µM BTB-(squares) induced ERK activation in CHO cells. **(D-F)** Förster-resonance energy transfer (FRET)-based EKAR biosensors were used to assess BTB-induced ERK activation in the **(D)** cytosol and **(E)** nucleus, and **(F)** its concentration dependence. **(G, H)** The requirement for GPR65 internalization in ERK activation was assessed in the **(G)** cytosol and **(H)** nucleus following pre-treatment with 30 µM PS2 or 0.1% (v/v) DMSO prior to stimulation with 100 µM BTB. Data represent means from four independent experiments. * *p-adj* < 0.05, ** *p-adj* < 0.01, **** *p-adj* < 0.0001; **(A,C)** two-way ANOVA (stimulation x condition); **(G-H)** two-way ANOVA (biosensor x condition), both with Bonferroni-corrected post hoc; **(B)** unpaired t-test.

Genetically encoded Förster resonance energy transfer (FRET) biosensors afford temporal resolution of signaling in live cells and they can be spatially restricted to quantify signaling in distinct subcellular compartments. ERK activity reporter (EKAR) FRET biosensors (*30*) were employed to spatiotemporally resolve GPR65-driven ERK signaling. Preliminary experiments revealed an inherent sensitivity of FRET biosensors to pH and thus experiments focused only on BTB-driven GPR65 signaling. BTB coordinated an increase in ERK activation in both the cytosol (Fig. 3D) and nucleus (Fig. 3E), with the cytosolic signal appearing more transient than the more persistent nuclear signal occurring across the 30-min measurement window. BTB-induced ERK activation in both compartments was concentration-dependent, with EC_50_ estimates of 40.3 µM for cytoEKAR and 0.77 µM for nucEKAR (Fig. 3F). To determine the contribution of endosomal GPR65 in driving compartment-specific ERK responses, further experiments were performed where mGPR65 cells expressing each biosensor were pre-treated with either DMSO or PS2, and the effect of 100 µM BTB (which gave robust responses in both compartments under basal conditions) was measured. There was a significant interaction between biosensors and experimental condition (*F*(1,12) = 36.666, *p* < 0.0001). Focusing on cytosolic ERK activity, a slight, but significant, increase was observed when endocytosis was inhibited, the signal persisting more towards the end of the recording period (DMSO, 0.756 ± 0.169, PS2, 0.977 ± 0.067, *p-adj =* 0.0265; Fig. 3G). Conversely, PS2-treatment completely abolished any ERK activity in the nucleus following challenge with BTB (DMSO, 0.716 ± 0.039, PS2, −0.206 ± 0.033, *p-adj* < 0.0001; Fig. 3H). As with endpoint ERK assays, PS2 treatment did not affect the ability of PDBu to coordinate ERK activity in either compartment (Condition x biosensor interaction: *F*(1,12) = 0.0607, *p* = 0.81; Fig. S3D). Data thus far highlight an ability of GPR65 agonists to induce receptor internalization, which subsequently affects the generation of signals that may be important for longer term readouts and functional properties. However, data so far have been collected in heterologous systems, with enhanced receptor expression and whether these also occur at the endogenous level remains to be explored.

### Disrupting GPCR internalization in primary mouse FLS affects signaling, transcriptional responses and the ability to coordinate sensory neuron sensitization

Within FLS, GPR65 activation has been shown to coordinate production of pro-inflammatory mediators capable of sensitizing dorsal root ganglia (DRG) sensory neurons, which may offer some explanation for the increased observance of pain-like behaviors in mice following activation of GPR65 within the knee-joint space (*11*). Accordingly, the role of receptor internalization in GPR65-driven FLS signaling was investigated. Studying the trafficking of endogenous GPCRs represents a challenge due to the lack of validated antibodies to localize GPCRs, thus an indirect approach of localizing clathrin within FLS following activation of GPR65 was employed to gauge whether GPR65 internalization occurs in primary cells. Within 10-min of stimulating FLS with pH 6.0, or 100 µM BTB a redistribution of clathrin to the plasma membrane was evident, an effect not seen for FLS which were stimulated with DMSO (Figs. 4A, B). Pre-treatment of FLS with the clathrin inhibitor PS2, reduced the generation of cAMP in response to GPR65 agonists (*F*(1,12) = 59.256, *p* < 0.0001; Fig. 4C), but did not affect basal cAMP levels, or the cAMP accumulation achieved in response to FSK stimulation (Condition Effect: *F*(1,12) = 1.0724, *p* = 0.321; Fig. S4A,B), providing further evidence that GPR65 likely internalizes and that this remains functionally important for signaling output in primary cells with endogenous receptor expression.

**Figure 4.**
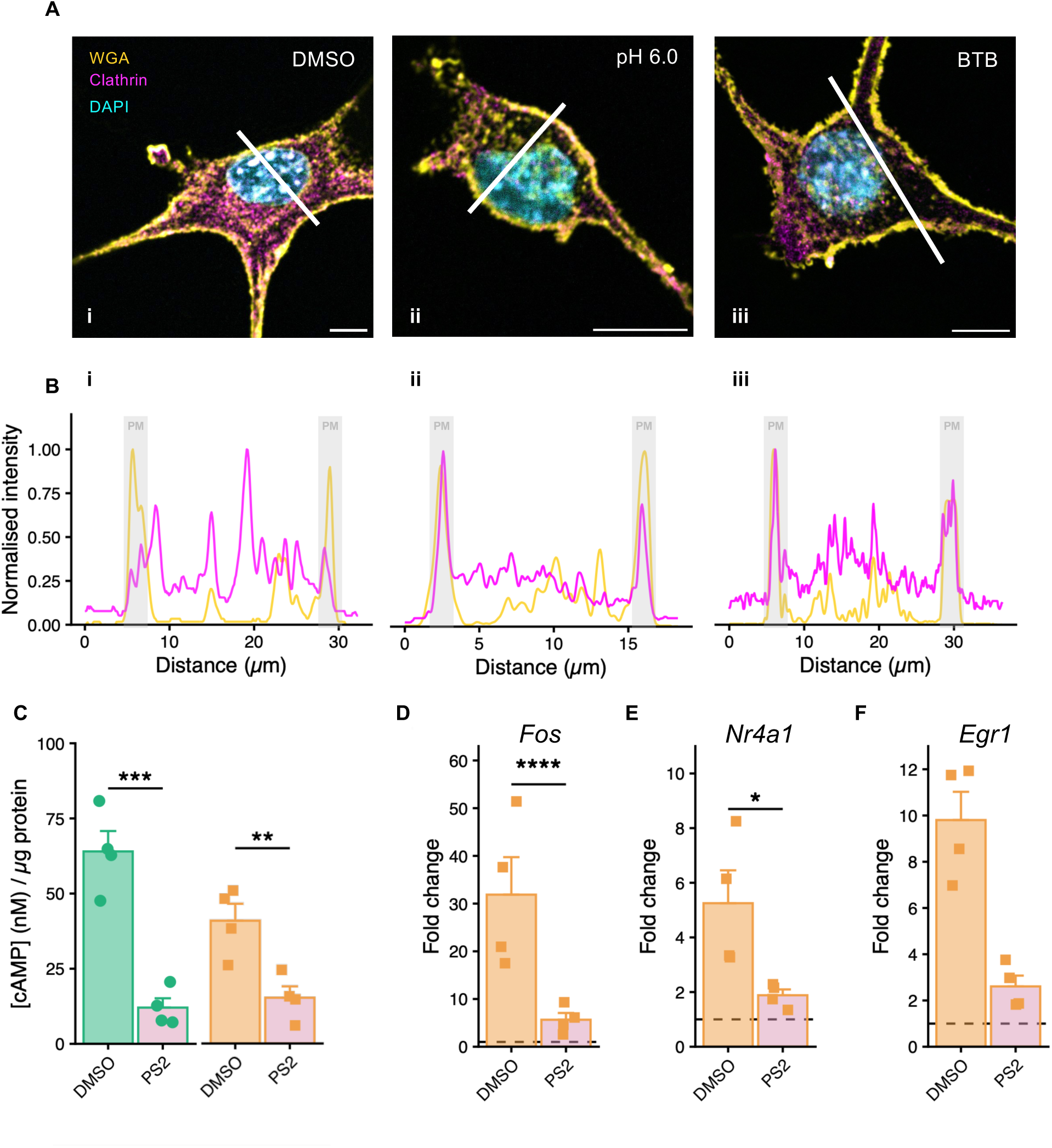
Disrupting GPR65 internalization alters signaling and transcriptional responses in FLS. **(A)** Representative staining of FLS following 10-min stimulation with **(i)** 0.1% (v/v) DMSO, **(ii)** HBSS (pH 6.0) or **(iii)** 100 µM BTB (pink, clathrin heavy chain; yellow, wheat germ agglutinin; blue, nuclear stain; scale-bar, 10 µm). **(B)** Intensity profiles of fluorescent signals across annotated transects (white line) from **(A)**. **(C)** Effect of receptor internalization on cAMP accumulation in FLS following pre-treatment with 50 µM PS2 or 0.1% (v/v) DMSO prior to stimulation with pH 6.0 (circles) or 100 µM BTB (squares). **(D-F)** Expression of GPCR-responsive immediate early genes **(D)** *Fos*, **(E)** *Nr4a1* and **(F)** *Egr1* following BTB stimulation, assessed by qPCR after pre-treatment with 50 µM PS2 or 0.1% (v/v) DMSO. Data in **(B)** are representative of a single transect from one cell; data in **(D-F)** represent means from four independent experiments. * *p-adj* < 0.05, ** *p-adj* < 0.01, *** *p-adj* < 0.001, **** *p-adj* < 0.0001; **(C)** two-way ANOVA (stimulation x condition); **(D-F)** three-way ANOVA (gene x stimulation x condition) performed on ΔCt values, all with Bonferroni-corrected post hoc tests.

Given that GPR65 activation on FLS can coordinate release of pro-inflammatory cytokines (*11*), the receptor likely engages transcriptional machinery upon activation. Since preventing GPR65 internalization reduces BTB-driven induction of a reporter gene (Fig. 3B) and nuclear EKR activation (Fig. 3H), the ability of BTB to coordinate expression of immediate-early genes commonly associated with GPCR signaling (*31*, *32*) was investigated using qPCR. A significant interaction between the abundance of mRNA for genes of interest and experimental condition was observed (*F*(2,36) = 6.614, *p* = 0.004), indicating that receptor internalization affects induction of immediate early genes. The stimuli cells received also had an effect (*F*(1,36) = 111.635, *p* < 0.0001), but further scrutiny revealed mRNA levels were consistent for vehicle treatment, according to each gene (*F*(1,36) = 0.0003, *p-adj* = 0.985), indicating a differing ability of BTB to induce immediate-early gene induction depending on whether internalization has been inhibited or not. PS2-pretreatment impaired the BTB-induction of *Fos* (ΔCts: DMSO, 9.894 ± 0.605, PS2, 13.3 ± 0.404, *p-adj <* 0.0001; Fig. 4D), and *Nr4a1* (DMSO, 17.297 ± 0.66, PS2, 18.601 ± 0.193, *p-adj =* 0.032; Fig. 4E). The induction of *Egr1,* however, did not significantly differ according to whether cells were pretreated with DMSO or PS2 (DMSO, 15.637 ± 0.531, PS2, 16.756 ± 0.122, *p-adj =* 0.0635; Fig. 4F). Interleukin-6 (IL-6) and monokine induced by gamma interferon (MIG) are among the inflammatory cytokines present in FLS media following BTB-simulation of GPR65 (*11*), thus, their corresponding genes, *Il6* and *Cxcl9* respectively, are likely to be induced by BTB in FLS. However, it was not possible to detect an induction of these genes in the DMSO-pretreated cells following the 90-min stimulation (*F*(1,12) = 0.16, *p* = 0.697; Fig. S4C), thus, this window is likely too short to detect the beginning of cytokine production. Experiments were not extended though due to the unknown potency of PS2 over longer durations.

Attempts to transfect FLS with an eGFP-expressing plasmid revealed poor transfection efficiency under various conditions and reagents, thus excluding use of DynK44E to study endosomal driven signaling of GPR65 in FLS. siRNA-mediated knockdown of clathrin was thus used as an alternative. Cells received two doses of siRNA targeting the heavy chain of clathrin (knockdown FLS), or a negative control siRNA (control FLS). 72-h post initial siRNA exposure, knockdown and control FLS were stimulated with 100 µM BTB and 24-h later media were collected. Media were also collected from FLS untreated with either siRNA or BTB (naive FLS; Fig. 5A). Immunoblotting of FLS lysates, prepared after media collection, revealed decreased expression of clathrin in knockdown FLS compared to control FLS, indicating successful siRNA-mediated knockdown (Fig. 5B and S5A). Naive lumbar DRG neurons isolated from mice, were cultured overnight in the conditioned media collected from FLS prior to electrophysiological characterization (Fig. 5C). Consistent with previous reports (*11*) conditioned media from BTB-treated control FLS reduced the rheobase of DRG sensory neurons compared to neurons incubated in the media taken from naive FLS. Furthermore, conditioned media from BTB-treated knockdown FLS failed to sensitize DRG sensory neurons, with excitability more comparable to cells incubated with naive FLS conditioned media (Control FLS, 146.3 ± 31.2 pA, naive FLS, 389.63 ± 69.4 pA, knockdown FLS, 379.1 ± 59.6 pA, *H*(2,87) = 9.416, *p* = 0.009; Fig. 5D). The effect of conditioned media on DRG sensory neuron activity was further explored by investigating the number of action potentials fired in response to 500 msec stimulations of increasing current, with the neurons that were incubated in conditioned media from control FLS stimulated with BTB being markedly more excitable (i.e., greater AP discharge) compared to those of other conditions (*F*(2,1232) = 20.138, *p* < 0.0001; Fig. 5E). Electrophysiological characterization revealed no other differences in the intrinsic or active properties of the DRG sensory neurons based on experimental condition (Table 1).

**Figure 5.**
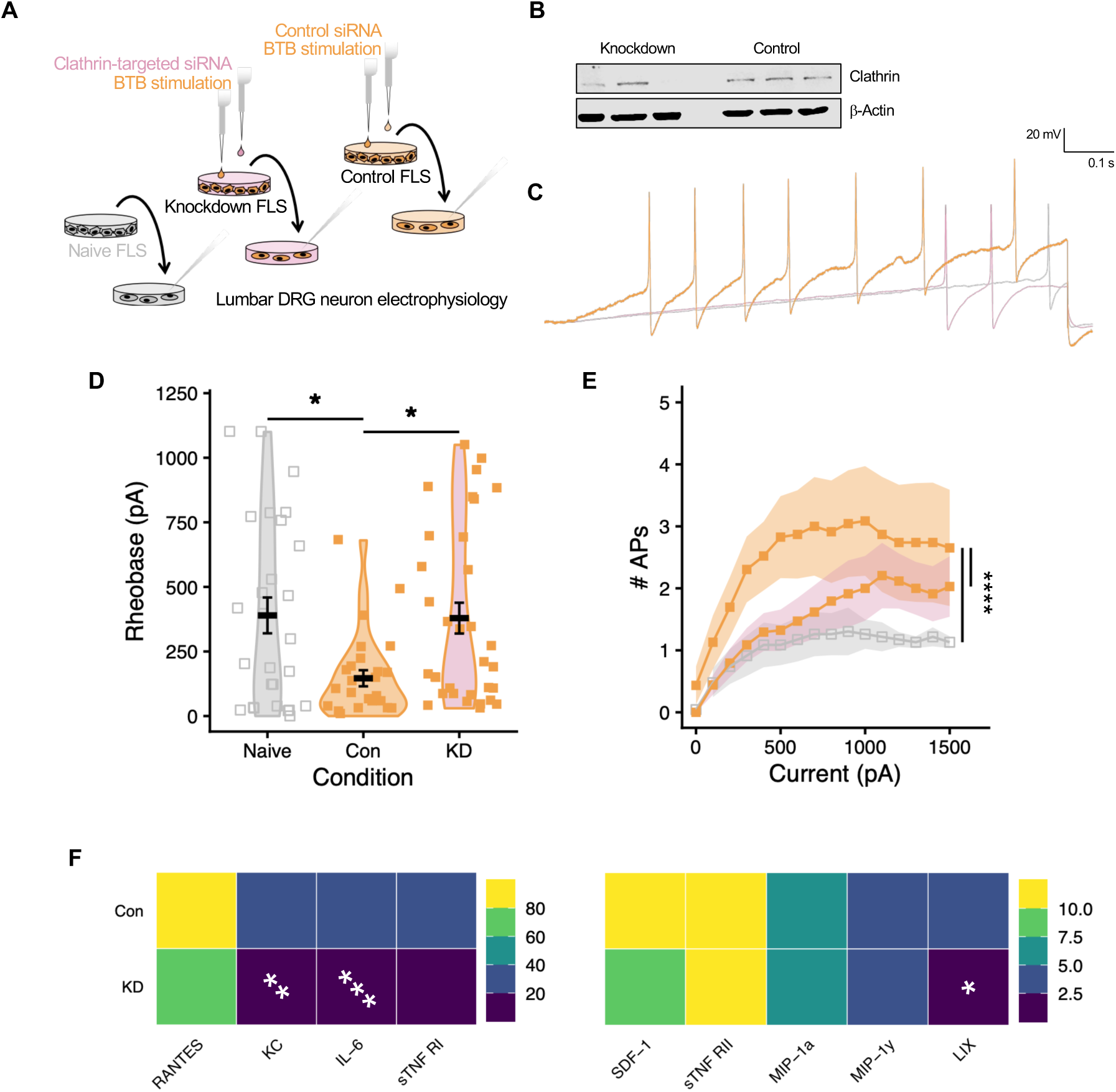
Reduced clathrin expression in FLS impairs their ability to sensitize DRG sensory neurons. **(A)** To impair receptor internalization over an extended period, FLS were transfected with siRNA targeting the clathrin heavy chain (knockdown; KD) or control (Con) siRNA. Conditioned media were collected 24 h after stimulation with 100 µM BTB; additional control media were collected from naive FLS. Media were applied to naive lumbar DRG neurons overnight. **(B)** Immunoblot confirmed reduced expression of clathrin heavy chain expression in siRNA-treated FLS. **(C)** Representative whole-cell electrophysiological recordings from lumbar DRG neurons following incubation with conditioned media during ramped current stimulation. **(D, E)** Neuronal excitability assessed by **(D)** rheobase and **(E)** input-output relationships. **(F)** Blot-based array quantification of inflammatory mediators in conditioned media from BTB-stimulated versus naive FLS. Data in **(C)** are representative traces from cells with similar capacitance; data in **(D)** represent individual cell measurements from four independent DRG neuron cultures; data in **(E)** represent means of cells recorded from four independent DRG neuron cultures; data in **(F)** represent four independent conditioned media collections pooled prior to analysis. * *p-adj* < 0.05, ** *p-adj* < 0.01, *** *p-adj* < 0.001, **** *p-adj <* 0.0001; **(D)** Kruskal-Wallis test (condition); **(E)** aligned rank transform ANOVA (current x condition); **(F)** two-way ANOVA (cytokine x condition), all with Bonferroni-corrected post hoc tests.

**Table 1.**
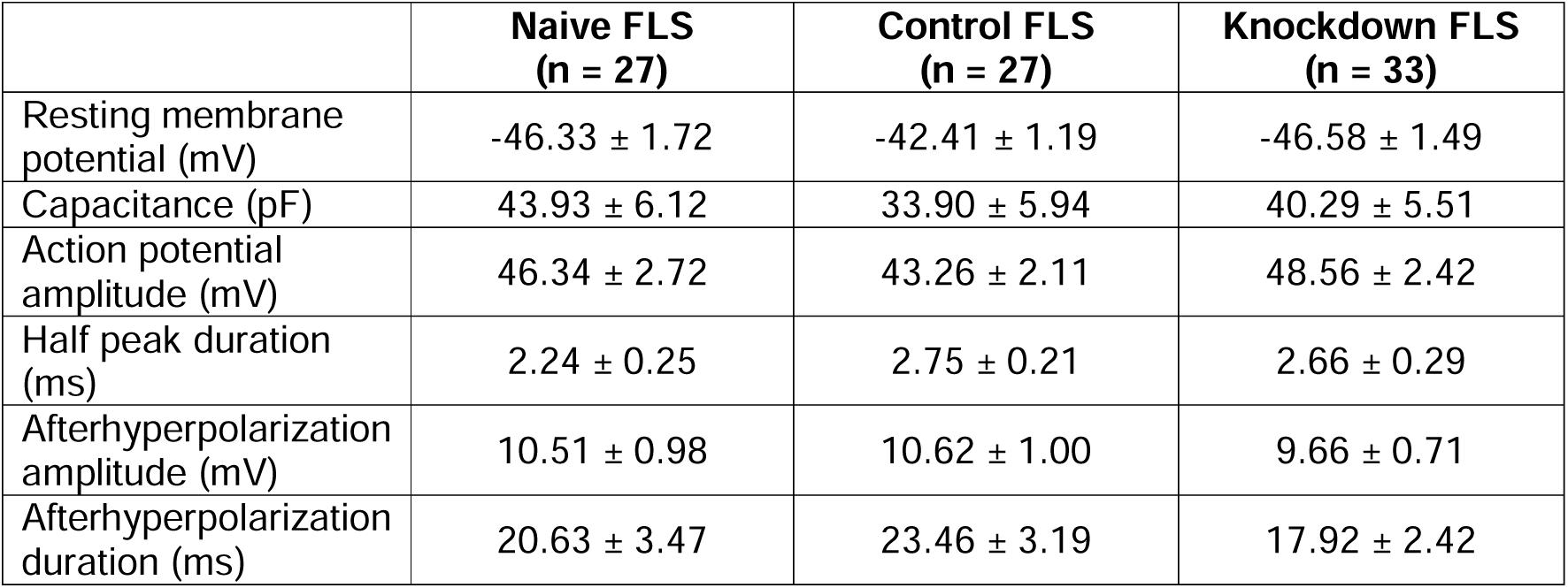
Intrinsic and active properties of lumbar dorsal root ganglion neurons (N = 4) cultured in conditioned media collected from FLS.

|  | Naive FLS<br>(n = 27) | Control FLS<br>(n = 27) | Knockdown FLS<br>(n = 33) |
| --- | --- | --- | --- |
| Resting membrane potential (mV) | -46.33 ± 1.72 | -42.41 ± 1.19 | -46.58 ± 1.49 |
| Capacitance (pF) | 43.93 ± 6.12 | 33.90 ± 5.94 | 40.29 ± 5.51 |
| Action potential amplitude (mV) | 46.34 ± 2.72 | 43.26 ± 2.11 | 48.56 ± 2.42 |
| Half peak duration (ms) | 2.24 ± 0.25 | 2.75 ± 0.21 | 2.66 ± 0.29 |
| Afterhyperpolarization amplitude (mV) | 10.51 ± 0.98 | 10.62 ± 1.00 | 9.66 ± 0.71 |
| Afterhyperpolarization duration (ms) | 20.63 ± 3.47 | 23.46 ± 3.19 | 17.92 ± 2.42 |

To begin to investigate the effects seen by conditioned media, an array-based approach was used to assay the presence of inflammatory mediators in samples, with differential mediator abundance observed between the samples (*F*(2,120) = 386.129, *p* < 0.0001; Fig. S5B). 9 cytokines were induced by BTB-stimulation of control FLS (relative to naive FLS), ordered greatest induction to lowest: RANTES, KC, IL-6, sTNF RI, SDF-1, sTNF RII, MIP-1α, MIP1-γ and LIX (Fig. 5F). Compared to control FLS, knockdown FLS produced significantly less KC (*p-adj* = 0.0034), IL-6 (*p-adj =* 0.0005) and LIX (*p-adj* = 0.0107). BTB stimulation of knockdown FLS significantly induced two cytokines – IL-7 and GCSF – relative to naive FLS, whereas control FLS showed no induction of these cytokines. Full analysis of cytokine presence is detailed in (Table S1). These data support the notion that BTB-induces internalization of GPR65 in FLS, which is necessary for coordinating pro-inflammatory responses, with the reduced levels of pro-inflammatory mediators produced by knockout FLS unable to sensitize sensory neurons.

## Discussion

Cellular communication is vital for coordination of the many specialized systems which support multicellular organisms; however, it can also drive disease. Activation of GPR65 on FLS in the knee joint triggers the release of inflammatory mediators capable of recruiting immune cells and sensitizing sensory neurons to drive inflammatory joint pain (*11*). The current study provides mechanistic detail between GPR65 activation on FLS and sensory neuron sensitization. In response to agonist stimulation of cells overexpressing a tagged version of the receptor, GPR65 internalized to early endosomes (Fig. 1) in a manner dependent on the actions of clathrin and dynamin (Fig. 2). Disrupting the function of endocytic machinery, and hence preventing GPR65 internalization, negatively affected the ability of protons and BTB to coordinate signaling responses in mGPR65 CHO cells, most notably induction of a CRE reporter and activation of ERK within the nucleus (Fig. 3). Stimulation of FLS with GPR65 agonists caused a redistribution of clathrin to the plasma membrane, indicative of similar agonist-induced internalization of endogenous GPR65 in this relevant primary cell type. Pre-treatment of FLS with the clathrin inhibitor PS2 also attenuated agonist-driven cAMP accumulation and induction of immediate-early genes (Fig. 4). siRNA-mediated knockdown of clathrin in FLS offered an insight into the long-term consequences of preventing receptor internalization. Following activation of GPR65 with BTB, reduced levels of inflammatory mediators were measured in the conditioned media of FLS with reduced clathrin expression, compared to control FLS. Consequently, the conditioned media from knockdown FLS failed to evoke sensitization of DRG sensory neuron activity (Fig. 5). This work thus highlights a necessity of GPR65 internalization for subsequent mediator production and communication with sensory neurons which results in their sensitization.

Although psychosine-induced trafficking of GPR65 was studied here (Figs. S1, S2), the consequences of receptor location and psychosine-evoked signaling output were not pursued, a decision owing to the distinct signaling signature of psychosine compared to protons and BTB (*11*). Investigation of endosomal signaling driven by this agonist however warrants further pursuit.

A study focusing only on proton-induced signaling of GPR65 has independently observed decreased cAMP accumulation following acidic challenge when dynamin is pharmacologically inhibited, strengthening the notion that subcellular localization of the receptor is a key determinant of signaling output. However, this study reports constitutive internalization of GPR65 and concludes that trafficking and compartmentalized signals are uncoupled (*33*). Data presented here do not unequivocally rule out the possibility that GPR65 passively shuttles between endomembrane compartments, but do suggest colocalization with early endosomes increases in response to agonist-treatment (Fig. 1). A key difference between these studies is the presence of intracellular GPR65 under basal conditions as assessed by microscopy. Our fluorescently tagged GPR65 is predominantly localized to the plasma membrane under basal conditions, or following exposure to vehicle (Figs. 1, S1), a contrast to Morales Rodríguez and colleagues’ approach to localize GPR65 via immunolabelling of FLAG-tag which can be seen intracellularly regardless of extracellular pH (*33*). Possible explanations for this include the differing cellular backgrounds and possible differences in receptor expression levels. Furthermore, antibody binding to GPCRs on live cells has been shown to promote internalization in its own right (*34*), which may offer explanation as to why FLAG-GPR65 was observed to be equally distributed among various endosomal compartments (*33*).

The findings reported here, implicate GPR65 localized within the endomembrane system as a determining feature of signaling output. This is not unusual for GPCRs, however, given the progressive acidification of the endomembrane system (*35*), and inherent sensitivity of GPR65 to protons, the influence of endosomal pH on GPR65 activity remains to be explored and raises questions about what drives continued activation of GPR65 following non-proton-induced internalization to early endosomes: endosomal acidity, ligand, or both? Furthermore, how cells remain sensitive to changes in extracellular pH fluctuation after GPR65 has been internalized would be interesting to study.

The experiments detailed here are reminiscent of peripheral sensitization, a major contributor to inflammatory pain. Although the experiments with FLS described here mainly utilize the exogenous GPR65 agonist, BTB (to avoid confounding activation of other proton-sensitive receptors by acid), it is hypothesized that the localized acidosis of inflammatory joint disorders such as arthritis (*11*, *36*, *37*) would activate GPR65 on FLS, trigger internalization and the induction of inflammatory cytokines which act to promote joint inflammation as well as sensitize neighboring nociceptive neurons and thus contribute to inflammatory joint pain in a disease setting. It is noteworthy that here IL-6, which was among the most upregulated cytokines detected following GPR65 activation of control FLS, was barely detected in the media of stimulated knockdown FLS (Fig. 5F), especially considering BTB stimulation of human FLS, also induces IL-6 (*11*) and the promising results of clinical investigation of an anti-IL-6 receptor monoclonal antibody (tocilizumab) in rheumatoid arthritis patients (*38*).

Here, pronociceptive effects of endosomal GPR65 are limited to sensitization of cultured DRG sensory neurons, it thus remains to be verified if subcellular GPR65 drives pain-like behavior *in vivo*. Intra-articular injection of BTB is sufficient to evoke a GPR65-driven inflammatory pain response (*11*) and thus administration of crude endocytic inhibitors, including PS2, may offer clarity, and such approaches have been used for other pronociceptive GPCRs (*19*, *21*). This would also allow assessment of whether GPR65 drives inflammation from endosomes, given two of the inflammatory mediators which were significantly blunted in release from knockdown FLS – KC and LIX (Fig. 5F) – are known to stimulate immune cell recruitment (*39*). Promising results, i.e. reductions in pain-like behaviors and inflammation following inhibition of endocytic machinery prior to GPR65 engagement, would specifically highlight subcellular GPR65 as a viable therapeutic target for achieving inflammation-specific analgesia, to address the inadequacies and risks of currently used medicines (*40*). Indeed, repurposing of a small-molecule GPR65 antagonist, which has recently been shown to reduce tumor growth and immune cell recruitment (*41*), for treatment of inflammatory pain might be worthwhile. Findings here would suggest strategies to increase delivery to endosomal compartments, i.e. lipid-conjugation (*42*) or nanoparticle-based delivery systems (*43*) might also enhance therapeutic benefit.

Overall, this work builds upon mounting evidence which suggests that the targeting of GPR65 may represent a means to achieve analgesia in inflammatory conditions and suggests that strategies targeting the receptor at endosomal platforms might enhance patient benefit. Furthermore, this work extends the notion of compartmentalized signaling as a driving force in nociception, to include the relevance on transcriptional output in a fully endogenous system. This may also be the first report of such a signaling paradigm occurring in fibroblast-like synoviocytes, opening further avenues for investigation in the study of cell-cell communication.

## Materials and Methods

### Animals

Use of animal tissues was regulated in accordance with the United Kingdom Animals (Scientific Procedures) Act 1986 Amendment Regulations 2012 and was approved by the University of Cambridge Animal Welfare Ethical Review Body (*44*). Tissues were collected from C57BL/6J wild-type mice (Envigo) of both sexes at age 10-12 weeks. Mice were housed in groups of up to five, with enrichments and *ad libitum* access to food and water in a holding room maintained at 21 °C and operating a 12-h light/dark cycle.

### Isolation and culture of mouse fibroblast-like synoviocytes

After humanely killing mice by cervical dislocation and exanguination, the knee joints were exposed by removing the skin, patellae were collected in phosphate-buffered saline after resecting the quadriceps muscles. Dissected patellae were then individually transferred to FLS media: Dulbecco’s Modified Eagle Medium/F-12 Nutrient Mixture + GlutaMAX (DMEM/F-12; Life Technologies) supplemented with 25% (v/v) fetal bovine serum (FBS; Merck) and 1% (v/v) penicillin/streptomycin (Life Technologies) in a 24-well plate and incubated at 37 °C, 5% CO_2_. Daily media changes were performed until a substantial number of FLS had grown out of the patellae, at which point the tissue was discarded and remaining adherent cells trypsinized; cells from the two patellae of each animal were combined in 6-well plates and maintained by refreshing media every other day. FLS were used for experiments from passage 3 (P3), at which point cultures are predominantly FLS (*45*) and cells were isolation matched for experiments (i.e. within each experiment each condition was assayed on cells from the same animal).

### Agonists and inhibitors

The GPR65 agonist, BTB09089 (BTB) was synthesized by Otava Chemicals and reconstituted at 100 mM in dimethyl sulfoxide (DMSO, Merck). 5 mM stocks of psychosine (Psy, Merck) were prepared in chrloroform:methanol (1:1). Forskolin (FSK, Merck), an adenylyl cyclase activator, was prepared at 10 mM in DMSO. The phorbol ester, phorbol 12,13-dibutyrate (PDBu, Tocris), an activator of mitogen-activated kinases, was prepared at 1 mM in DMSO. Coelenterazine H (Nanolight Technologies), a substrate of the Renilla luciferase, was prepared at 500 µM in 100% methanol. The clathrin inhibitor, PitStop2 (PS2; Abcam) was prepared at 30 mM in DMSO. Where used, all stock concentrations were diluted in neutral pH assay buffers (pH 7.4).

### siRNA knockdown of clathrin

Once confluent, P3 FLS, plated in 6-well plates, were incubated with 30 pmol of either: two siRNAs targeting the clathrin heavy chain gene (*Cltc*, Assays 220124 and 220125; Product # AM16708, Invitrogen), or a negative control siRNA (AM4611; Invitrogen). siRNAs were diluted in OptiMEM (Life Technologies) with Lipofectamine RNAiMax reagent (Invitrogen), according to the manufacturer’s guidelines. Cells received two doses of siRNAs, with a 24-h gap between transfections. 24-h after the second transfection cells underwent serum starvation (unsupplemented DMEM/F-12 media) for 6-h, before stimulation with 100 µM BTB. 24-h post stimulation, media was collected and immediately stored at −80 °C. Cells were then washed with ice-cold PBS, collected in 500 µl lysis buffer (50 mM Tris-HCl (pH 8.0), 150 mM NaCl, 1% (v/v) Triton X-100) and agitated on ice for 10-min before centrifugation (13,200 rpm, 15-min, 4 °C). The supernatant was collected and protein content quantified via Bradford assay before combination with 2X laemmli buffer (125 mM Tris-HCl (pH 6.8), 20% (v/v) glycerol, 10% (v/v) β-mercaptoethanol, 4% (w/v) sodium dodecyl sulfate, 0.005% (w/v) bromophenol blue) and stored at −20 °C.

### Western blotting

After 5-min heating at 70 °C, 50 µg of each lysate was loaded on an 8% SDS PAGE gel and resolved by electrophoresis (20-min 60V, 150-min 110V). NB – siRNA transfections were conducted four times independently; however, sufficient protein was only extracted from three of each of the conditions (knockdown or control) to load 50 µg; blotting with 30 µg did not result in sufficient transfer to detect clathrin. Wet transfer to PVDF membrane (Amersham) was conducted overnight (0.1A, 16-h) within a cold room. Transfer buffer contained 10% (v/v) methanol and 0.05% (w/v) sodium dodecyl sulfate. The membrane was blocked in 5% (w/v) bovine serum albumin (BSA; Merck) prepared in tris-buffered saline containing 0.1% (v/v) Tween 20 (TBST; 6-h, 4 °C) before the addition of primary antibodies raised against clathrin heavy chain (both 1:1,000, rabbit monoclonal, Cell Signaling Technology, 4796 and mouse monoclonal, Invitrogen, MA1-065) and β-actin (1:1,000, goat polyclonal, A121645, Antibodies.com) and further overnight incubation (4 °C). Following washing with TBST, the membrane was incubated with secondary antibodies: anti-rabbit IgG-IRDye 680RD (1:10,000, donkey host, LI-COR, 926-680073), anti-mouse IgG-IRDye 680RD (1:10,000, donkey host, LI-COR, 926-68072), and anti-goat IgG-IRDye 800CW (1:10,000, donkey host, LI-COR, 926-32214) for 1-h at room temperature, before washing and imaging with an Odyssey XF imager (LI-COR).

### Immunocytochemistry

P3 FLS were plated on poly-D-lysine coated glass coverslips and cultured for 24-h. Cells were washed twice before a 10-min incubation (37 °C) with assay buffer (HBSS) containing 10 mM HEPES (Merck), 10 mM MES (Fisher Scientific) and 2 mg/ml BSA, pH 7.4, herein referred to as HBSS buffer), containing 0.1% (v/v) DMSO, 100 µM BTB or buffered to pH 6.0. After which cells were washed once with PBS before being fixed with 4% (w/v) paraformaldehyde (PFA; Merck) for 20-min at room temperature. After 2 washes with PBS containing 0.001% (v/v) Tween 20 (PBS-T; Merck), cells were incubated with 1 µg/ml wheat germ agglutinin conjugated with CF568 (20977-1, Biotium) for 10-min at room temperature, after which cells were washed twice with PBS-T. Cells were then incubated with antibody diluent (PBS containing 0.02% (v/v) Triton X-100 (Merck), 5% (v/v) donkey serum (Merck) and 1% (w/v) BSA) for 1-h to block and permeabilize before overnight incubation with primary antibody: anti-clathrin heavy chain (1:100, rabbit polyclonal, Cell Signaling Technology, 4796) at 4 °C. The following day cells were washed thrice with PBS-T before incubation with anti-rabbit IgG-AF488 (1:1,000, donkey polyclonal, Invitrogen, A32790) for 2-h at room temperature. Cells were washed once with PBS-T before 5-min incubation with 300 nM DAPI dihydrochloride (Invitrogen), followed by a further two PBS-T washes and finally mounting on glass slides with Mowiol mounting media (6.66% (w/v) Mowiol 4-88 (Merck) and 0.2% (w/v) DABCO (Merck)). Slides were imaged with a Leica SP5 laser-scanning confocal microscope and 63x oil objective. Profile analyses were performed in Fiji (*46*), fluorescence intensities across each transect were normalized to maximal intensity per transect, per channel.

### cAMP assays (FLS)

P3 FLS were plated in standard 24-well plates, once confluent cells were washed twice and incubated with HBSS buffer containing 50 µM PS2 or 0.1% (v/v) DMSO for 30-min (37 °C). At the end of the incubation assay buffer was replaced with 400 µl fresh assay buffer or assay buffer containing 100 µM BTB, 10 µM FSK, or buffered to pH 6.0 and cells returned to the incubator. After 15-min of stimulation, plates were transferred to ice, stimulatory solutions aspirated and 50 µl lysis buffer (50 mM HEPES, 10 mM CaCl_2_, 0.35% (v/v) Triton X-100, pH 7.4) was added to each well, the plate was incubated on ice with gentle agitation for 30-min. The concentration of cAMP in lysates was quantified using LANCE Ultra cAMP detection reagents (Revvity), each lysate was assayed in duplicate alongside a cAMP standard curve. To account for differences in the cell number between conditions the total protein content of lysates was also assayed using Bradford reagent (Merck) and a standard curve made up of known amounts of BSA. The cAMP in lysates was interpolated from the standard curve and normalized to total protein content.

### qPCR

Once confluent, P3 FLS in 6-well plates, were serum-starved overnight with unsupplemented DMEM/F12. The following day cells were washed twice before a 30-min incubation with either 50 µM PS2 or 0.1% (v/v) DMSO made up in HBSS buffer. After pre-incubations, cells were exposed to HBSS buffer containing DMSO or PS2, with half of each set of cells also exposed to 100 µM BTB. Cells were returned to the incubator for 90-min, after which stimulatory solutions were aspirated, and cells were lysed with TRIzol reagent (Merck) for RNA purification. After quantification RNA was immediately stored at −80 °C. cDNA was prepared once all RNA samples had been collected using high-capacity reverse transcription reagents (Thermo Fisher) and 500 ng RNA as template. Control reactions, where reverse transcriptase was omitted from the reaction mixture, were set up in parallel. Gene expression was assayed using TaqMan assays (Thermo Fisher) and a StepOnePlus System (Thermo Fisher) as per the manufacturer’s guidelines. Each reaction contained 2.5 ng of the relevant cDNA sample and each sample was assayed in duplicate per gene of interest. Negative controls, using the mock reverse transcription reaction as template, run in parallel exhibited no amplification. The probes used were: *Fos* (Mm00487425_m1), *Nr4a1* (Mm01300401_m1), *Egr1* (Mm00656724_m1), *IL-6* (Mm00446190_m1), *Cxcl9* (Mm00434946_m1) and *18SrRNA* (Mm03928990_g1). The average Ct of each sample/gene combination was calculated, ΔCt was calculated as Ct_Gene_-Ct*_18srRNA_*, the BTB-induced induction of expression was calculated by ΔCt_BTB-treated_ – ΔCt_Veh.-treated_ yielding ΔΔCt for each sample pairing and the fold change in expression was calculated as 2^-ΔΔCt^. Statistical analysis was performed on ΔCt values.

### Antibody array for inflammatory cytokine release

Condition media were analyzed using a Mouse Inflammation Antibody Array (Abcam), following the manufacturer’s guidelines. A total of 2 ml, pooled from four separate biological replicates, was incubated with membranes overnight at 4 °C. Chemiluminescent signals were captured with an Odyssey XF imager (LI-COR). Densitometry analysis was performed in Fiji (*46*): for each array the intensities of targets were normalized to the average intensity of positive control spots, yielding relative densities. Fold-change was calculated as the relative density of test conditions / relative density for the naive sample. Statistical analysis was performed on relative densities.

### Isolation and culture of mouse dorsal root ganglia (DRG) neurons

Mice were sacrificed by cervical dislocation before lumbar DRG (L2-L5) were collected in dissociation medium (L-15 + GlutaMAX growth media supplemented with 24 mM NaHCO_3;_ Life Technologies); only L2-L5 were collected because these DRG richly innervate the knee joint (*47*) and thus are most relevant to studying FLS – DRG neuron interactions. DRG were then incubated in dissociation media containing 1 mg/ml type 1A collagenase (Merck) and 6 mg/ml BSA for 15-min at 37 °C, 5% CO_2,_ before a further 30-min incubation in dissociation media containing 1 mg/ml trypsin (Merck) and 6 mg/ml BSA. DRGs were then suspended in culture media (L-15 + GlutaMAX growth media supplemented with 10% (v/v) FBS, 24 mM NaHCO_3_ and 2% (v/v) penicillin/streptomycin before several rounds of gentle mechanical titration and brief centrifugation. After sufficient trituration, dissociated cells were pelleted, resuspended in culture media and plated on glass-bottomed MatTek dishes. Cells were allowed to adhere for 2-h (37 °C, 5% CO_2_) before the addition of 200 µl conditioned FLS media and return to the incubator overnight. Conditioned medias from all experimental conditions were tested on each DRG preparation, after addition of conditioned medias, the experimenter was blinded to experimental conditions. Dishes were incubated within a 15 cm dish with a separate culture dish containing only 2 ml PBS to maintain humidity and prevent DRG cultures drying out during this low-volume culture.

### DRG neuron electrophysiology

Electrophysiology recordings were performed the day after cultures were prepared, using an EPC-10 amplifier (HEKA) and corresponding Patchmaster software. The extracellular solution contained (in mM): NaCl (140), KCl (4), MgCl_2_ (1), CaCl_2_ (2), glucose (4) and HEPES (10), pH 7.4. Patch pipettes were pulled from borosilicate glass capillaries (Hilgenberg) using a P-97 pipette puller (Sutter Instruments) with resistances of 3-6 MΩ and back-filled with intracellular solution containing (in mM): KCl (110), NaCl (10), MgCl_2_ (1), egtazic acid (EGTA, 1), HEPES (10), Na_2_ATP (2) and Na_2_GTP (0.5), pH 7.3. Whole cell currents were acquired at 20 kHz. Stepwise depolarization (Δ10 pA, 50 ms) was used to determine action potential threshold (rheobase) of cells. Only cells which fired action potentials were included in analyses. Action potential parameters were measured using Fitmaster (HEKA) and IgorPro (Wavemetrics) software. The excitability of neurons was further assessed by subjecting cells to longer stepwise increments of current injection (0 – 1,500 pA, Δ100 pA, 500 ms) and recording the number of action potentials discharged during stimulation.

### Cell line culture

All cell lines were cultured at 37 °C in a humidified 5% CO_2_ incubator. Parental Flp-In CHO cells (Thermo Fisher) were maintained in F12 media (Life Technologies) supplemented with 10% (v/v) FBS, 1% (v/v) penicillin/streptomycin and 100 µg/ml zeocin (InvivoGen). CHO cells stably expressing mouse GPR65 with N-terminal FLAG- and C-terminal -HA tags have previously been described (*11*) and were cultured in F12 media supplemented with 10% (v/v) FBS, 1% (v/v) penicillin/streptomycin and 550 µg/ml hygromycin-B (InvivoGen).

### cDNAs

A plasmid encoding mGPR65 tagged with the fluorescent mCherry at the C-terminus was created by standard subcloning practices using a LAMP1-mCherry vector kindly provided by Dr David Prole (University of Cambridge) (*48*), the construct was verified by Sanger sequencing. mGPR65-RLuc8 has previously been described (*11*). Plasmids encoding Rab5a-Venus as well as the wild-type (DynWT) and mutant dynamin (DynK44E) were provided by Profs. Nigel W. Bunnett (New York University) and Meritxell Canals (University of Nottingham) (*19*). A plasmid encoding enhanced green fluorescent protein (eGFP) was sourced from Clontech. A CRE-driven reporter plasmid (CRE-NLuc) was provided by Prof Graham Ladds (University of Cambridge) (*29*). Plasmids encoding FRET biosensors were sourced from AddGene: CytoEKAR (18679) and NucEKAR (18681) (*30*).

### Confocal microscopy

Flp-In CHO cells were plated in 6-well plates, once 70% confluent cells were transfected with 500 ng mGPR65-mCherry using Lipofectamine LTX (Invitrogen) at a ratio of 1:3 cDNA:LTX. 24-h post transfection cells were seeded onto poly-D-lysine coated glass coverslips and cultured overnight. The following day cells were washed twice and allowed to acclimatize in HBSS buffer for 30-min, after which cells were stimulated for 30-min at 37 °C with HBSS (pH 6.0), 100 µM BTB or 1 µM psychosine, or with HBSS (pH 7.4), 0.1% (v/v) DMSO or 0.02% (v/v) chloroform:methanol (1:1) as the respective vehicle controls. After stimulations, solutions were aspirated, cells were washed once with ice cold PBS before fixation with ice cold methanol for 10-min. After a series of PBS washes, cells were incubated with 300 nM DAPI dihydrochloride, washed twice more with PBS-T and finally coverslips were mounted onto microscopy slides with Mowiol mounting media. Cells were imaged with a Lecia SP5 confocal microscope, using the 68x oil objective lens, the same acquisition parameters were used to collect images of at least 5 individual cells from 3 separate transfections/stimulations. The number of intracellular mCherry puncta was quantified using Fiji (*46*): briefly, a Gaussian blur of 2 pixels was applied, before 0.1% contrast enhancement. The intracellular region was identified using the halo-fluorescence of mCherry at the cell membrane, the area of this region was measured, and intracellular puncta were identified using the Find Maxima algorithm. The number of puncta was then normalized to intracellular area.

### BRET assays

Flp-In CHO cells were plated in 10 cm dishes, once 70% confluent, cells were transiently transfected with 1 µg mGPR65-RLuc8 and 4 µg Rab5a-Venus or pcDNA3.1 using lipofectamine LTX reagent at a ratio of 1:3, cDNA:LTX. For experiments assessing the role of dynamin in receptor internalization, cells were transfected in 6-well plates, receiving 1 µg relevant dynamin construct or pcDNA3.1, 200 ng mGPR65-RLuc8 and 800 ng Rab5a-Venus, control cells received 1.8 µg pcDNA3.1 and 200 ng mGPR65-RLuc8. For experiments with PS2, cells were also transfected in 6-well plates, receiving 1.6 µg Rab5a-Venus or pcDNA3.1 and 400 ng mGPR65-RLuc8. 24-h post-transfections, cells were seeded on white opti-96well plates at a density of ~30,000 cells/well. The following day culture media was removed, and cells were washed twice before a 30-min incubation in HBSS buffer. For studies assessing clathrin-dependency cells were incubated with either 30 µM PS2 or 0.1% (v/v) DMSO. After initial acclimatization, cells were stimulated with 100 µl HBSS containing 5 µM coelenterazine H and various concentrations of agonist or pH buffered as necessary. Dual luminescence was quantified using a CLARIOstar (BMG Labtech) with emission sequentially recorded at 475 ± 30 nm and 535 ± 30 nm. Cells were maintained at 37 °C throughout measurements and each condition was assayed in duplicate. The ratio of Venus emission to RLuc8 emission constitutes the BRET ratio, netBRET was calculated by subtracting the BRET ratios from equivalent stimulation of cells expressing pcDNA3.1 in place of Rab5a-Venus, these were vehicle corrected to yield ΔnetBRET.

### End-point signaling assays (cell lines)

mGPR65-CHO cells were plated in 6-well plates, once 70% confluent cells were transfected with 1 µg pcDNA3.1 or Dynamin-K44E, 75 ng eGFP was co-transfected to visually confirm robust transfection. 24-h post transfection cells were seeded into standard 96-well plates at a density of 20,000 cells/well. For pERK assays cells underwent overnight serum starvation (OptiMEM). Before stimulations, cells were acclimatized in HBSS buffer for 30-min, 37 °C. Cells were stimulated with vehicle, 100 µM BTB or HBSS (pH 6.0), positive controls were included for each experiment: 10 µM FSK for cAMP and 1 µM PDBu for pERK. After 15-min of stimulation (37 °C) plates were immediately transferred to ice, stimulatory solutions removed and 50 µl lysis buffer (50 mM HEPES, 10 mM CaCl_2_, 0.35% (v/v) Triton X-100, pH 7.4) added, cells were agitated for 30-min at room temperature to ensure full lysis. cAMP or phosphorylated ERK were quantified in lysates using Lance Ultra TR-FRET reagents from Revvity as per the manufacturer’s guidelines, TR-FRET was read on a CLARIOstar PLUS (BMG Labtech) and normalized to that achieved by the positive control of each condition.

### EKAR sensor FRET assays

mGPR65 CHO cells were plated in 10 cm dishes, once 70% confluent cells were transfected with 2.5 µg cytoEKAR or nucEKAR CFP/YFP FRET biosensor. 24-h post-transfection cells were plated in black 96-well plates at a density of 30,000 cells/well, 8-h later media was replaced with OptiMEM for overnight serum starve. The following day cells were washed twice and allowed to acclimatize in 90 µl HBSS buffer for 30-min. For experiments assessing the role of internalization in EKAR activation, cells were preincubated with either 30 µM Pitstop2 or 0.1% (v/v) DMSO. After the 30-min pre-incubation, plates were moved to a CLARIOstar (BMG Labtech) set at 37 °C. Wells were excited at 425 ± 10 nm, with emission sequentially measured at 490 ± 20 nm and 550 ± 50 nm every minute. A baseline of 4-min was recorded before cells were challenged with 10 µl 10X concentrations of BTB or PDBu (1 µM final) as a positive control, measurements continued for a further 30-min. ERK activity was inferred from the ratio of YFP to CFP, which was vehicle and baseline corrected to yield ΔFRET.

### Reporter gene assays

CHO cells (70% confluent, 6-well plate) were transiently transfected with 500 ng CRE-NLuc, 500 ng mGRP65-mCherry and 1 µg either pcDNA3.1 or Dynamin K44E, 24-h later, cells were seeded at 30,000 cells/well in optiMEM. Cells for reporter gene analysis were plated in standard 96-well plates (the assay plate), additional wells were also seeded in black optiplates. The mCherry intensity of the wells seeded in black plates was measured the following day (Ex: 570 ± 25 nm; Em: 620 ± 20 nm) using a CLARIOstar (BMG Labtech) to ensure comparable transfection efficiencies between conditions. The day after plating, 2X BTB (100 µM final) prepared in optiMEM was added to wells of the assay plate, a subset of wells received an equivalent concentration of DMSO. Plates were incubated for 6-h (37 °C), at which point cells were lysed with Nano-Glo Luciferase Assay System reagents containing the luciferase substrate (Promega). Plates were protected from ambient light, gently agitated for 5-min at room temperature, before luciferase activity was measured on a CLARIOstar (BMG Labtech). The relative luminescent intensity of BTB-treated wells was divided by that of DMSO-treated wells to infer BTB-induction of NLuc expression for each condition.

### Data analysis and statistics

Data are presented as mean ± standard error of the mean (SEM), the number of experimental replicates is detailed in individual figure legends. Appropriate analyses were selected according to the number of factors being compared and whether data met the assumptions for parametric analyses, the statistical tests employed are stated in corresponding figure legends. All analyses were performed in R, and *P* values < 0.05 were considered significant.

## Supplementary Materials

This PDF file includes:

Figs. S1-S5.

Table S1.

## Acknowledgements

We thank Prof. Nigel W. Bunnett (New York University), Prof. Meritxell Canals (University of Nottingham), Prof. Graham Ladds (University of Cambridge) and Dr David Prole (University of Cambridge) for sharing plasmids necessary for experiments described. The mouse monoclonal anti-clathrin heavy chain primary antibody was a gift from Dr James Edgar (University of Cambridge).

## Author contributions

L.A.P. conceived and designed the project. L.A.P. and M.D. performed experiments. L.A.P. analyzed data. L.A.P. and E.S.S. secured funding. L.A.P. wrote the manuscript. All authors revised the manuscript.

## Data availability

All data sets supporting the conclusions of the paper are available in the University of Cambridge Apollo Repository (DOI: 10.17863/CAM.130775). For the purpose of open access, the authors have applied a Creative Commons Attribution (CC BY) license to any Author Accepted Manuscript version arising from this submission.

