## Supplemental Materials for "Endosomal GPR65 signaling in fibroblast-like synoviocytes promotes inflammatory cytokine release and nociceptive neuron sensitization"

**Supplementary Figure 1**

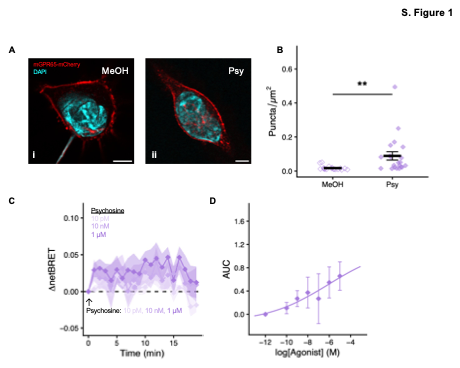

**Figure S1. Characterization of psychosine-induced internalization of GPR65. (A)** Representative images of Flp-In CHO cells expressing GPR65-mCherry following 30-min stimulation with **(i)** 0.02% (v/v) methanol:chloroform or **(ii)** 1 µM psychosine (red, GPR65-mCherry; blue: nuclear stain; scale bar, 5 µm). **(B)** Quantification of intracellular fluorescent puncta normalized to total cell area. GPR65 internalization following psychosine stimulation was assessed by BRET between GPR65-RLuc8 and Rab5a-Venus: **(C)**: kinetics and **(D)** concentration-response curve. Data in **(B)** represent individual cells from three independent experiments; data in **(C, D)** represent means from four independent experiments. ** *p-adj* < 0.01; Kruskal-Wallis test (stimulation) with Bonferroni-corrected post hoc.

**Supplementary Figure 2**

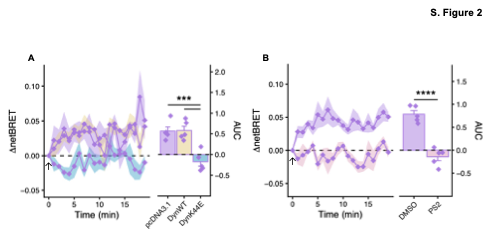

**Figure S2. Dynamin- and clathrin-dependence of psychosine-induced GPR65 internalization.** Trafficking of GPR65 to early endosomes in transfected Flp-In CHO cells following treatment with 1 µM psychosine was assessed by BRET between GPR65-RLuc8 and Rab5a-Venus. **(A)** Cells were transfected with wild-type dynamin (DynWT), a dominant negative mutant (DynK44E) or pcDNA3.1. **(B)** Cells were pretreated with the clathrin inhibitor PS2. Data represent five independent experiments. *** *p-adj* < 0.001, **** *p-adj* < 0.0001; two-way ANOVA (stimulation x condition) with Bonferroni-corrected post hoc.

**Supplementary Figure 3**

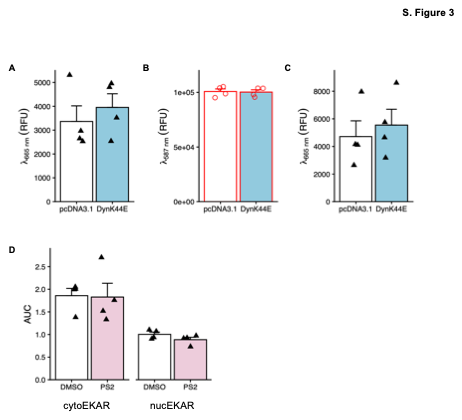

**Figure S3. Signaling assay controls for cell line experiments.** **(A)** Acceptor intensity of TR-FRET reagents used to measure cAMP in response to FSK stimulation was not affected by DynK44E overexpression. **(B)** Comparable mCherry intensity was observed across reporter assay conditions, indicating similar transfection efficiency. **(C)** Acceptor intensity of TR-FRET reagents used to measure ERK activation in response to PDBu stimulation was not affected by DynK44E overexpression. **(D)** Pretreatment with PS2 did not affect the FRET signal (i.e., ERK activation) induced by PDBu in cells expressing cytoEKAR or nucEKAR biosensors. Data represent means from four independent experiments.

**Supplementary Figure 4**

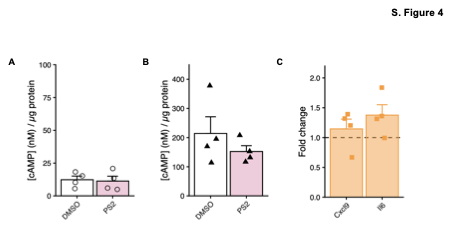

**Figure S4. FLS signaling assay controls and inflammatory mediator expression.** **(A, B)** The concentration of cAMP accumulated in response to **(A)** vehicle and **(B)** FSK treatment was not affected by PS2 pretreatment. **(C)** BTB-induced expression of inflammatory mediators (*Cxcl9***,** chemokine (C-X-C motif) ligand 9; *Il6,* interleukin-6) after 90-min stimulation under basal conditions. Data represent means from four independent experiments.

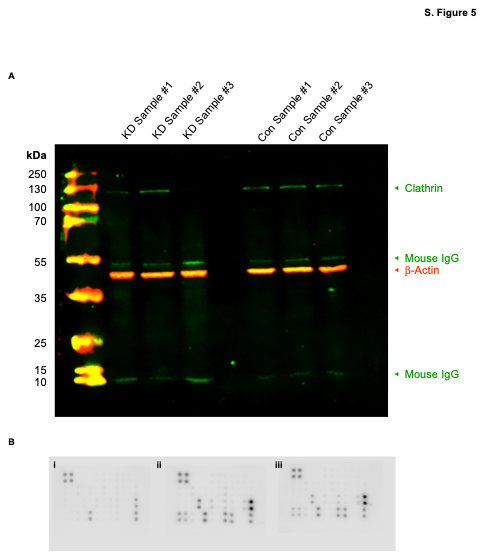
**Supplementary Figure 5**

**Figure S5. Raw blot images.** **(A)** Uncropped blot confirming siRNA-mediated knockdown of clathrin heavy chain in FLS. Inputs (L-R): PageRuler Plus Prestained Protein Ladder (10 to 250 kDa), targeted siRNA-treated samples #1-3, empty lane, control siRNAtreated samples #1-3. Red, IRDye 800 conjugated anti-goat-IgG staining. Green, IRDye 680 conjugated anti-mouse-IgG and anti-rabbit-IgG staining. **(B)** Images of array membranes used to assess inflammatory mediators in conditioned media from **(i)** naïve FLS, **(ii)** control FLS stimulated with BTB, **(iii)** knockdown FLS stimulated with BTB.

**Table S1.** Comparison of inflammatory cytokine levels in the conditioned media collected from mouse FLS after 24 hours culture in regular media (Control), DMSO or BTB. * p-adj < 0.05, ** p-adj < 0.01: Two-way ANOVA followed by Bonferroni-corrected posthoc.

| **Cytokine** | **Condition 1** | **Condition 2** | **p-adj** | **p-adj sifnic.** |
| --- | --- | --- | --- | --- |
| BLC | Naive | Con | 0.324000 | ns |
| BLC | Naive | KD | 0.062200 | ns |
| BLC | Con | KD | 0.342000 | ns |
| CD30 L | Naive | Con | 1.000000 | ns |
| CD30 L | Naive | KD | 0.431000 | ns |
| CD30 L | Con | KD | 0.253000 | ns |
| Eotaxin | Naive | Con | 1.000000 | ns |
| Eotaxin | Naive | KD | 0.810000 | ns |
| Eotaxin | Con | KD | 1.000000 | ns |
| Eotaxin-2 | Naive | Con | 0.577000 | ns |
| Eotaxin-2 | Naive | KD | 0.843000 | ns |
| Eotaxin-2 | Con | KD | 1.000000 | ns |
| Fas Ligand | Naive | Con | 1.000000 | ns |
| Fas Ligand | Naive | KD | 1.000000 | ns |
| Fas Ligand | Con | KD | 0.800000 | ns |
| Fractalkine | Naive | Con | 0.302000 | ns |
| Fractalkine | Naive | KD | 0.669000 | ns |
| Fractalkine | Con | KD | 1.000000 | ns |
| GCSF | Naive | Con | 0.011600 | * |
| GCSF | Naive | KD | 0.005550 | ** |
| GCSF | Con | KD | 0.001010 | ** |
| GM-CSF | Naive | Con | 0.974000 | ns |
| GM-CSF | Naive | KD | 1.000000 | ns |
| GM-CSF | Con | KD | 1.000000 | ns |
| I-TAC | Naive | Con | 0.181000 | ns |
| I-TAC | Naive | KD | 1.000000 | ns |
| I-TAC | Con | KD | 0.406000 | ns |
| IFNγ | Naive | Con | 1.000000 | ns |
| IFNγ | Naive | KD | 1.000000 | ns |
| IFNγ | Con | KD | 1.000000 | ns |
| IL-10 | Naive | Con | 1.000000 | ns |
| IL-10 | Naive | KD | 1.000000 | ns |
| IL-10 | Con | KD | 1.000000 | ns |
| IL-12 p40/p70 | Naive | Con | 0.060200 | ns |
| IL-12 p40/p70 | Naive | KD | 0.031300 | * |
| IL-12 p40/p70 | Con | KD | 0.927000 | ns |
| IL-12 p70 | Naive | Con | 1.000000 | ns |
| IL-12 p70 | Naive | KD | 1.000000 | ns |
| IL-12 p70 | Con | KD | 1.000000 | ns |
| IL-13 | Naive | Con | 1.000000 | ns |
| IL-13 | Naive | KD | 0.428000 | ns |
| IL-13 | Con | KD | 0.956000 | ns |
| IL-17 | Naive | Con | 1.000000 | ns |
| IL-17 | Naive | KD | 1.000000 | ns |
| IL-17 | Con | KD | 0.926000 | ns |
| IL-1β | Naive | Con | 0.053200 | ns |
| IL-1β | Naive | KD | 0.053200 | ns |
| IL-1β | Con | KD | 1.000000 | ns |
| IL-1α | Naive | Con | 0.382000 | ns |
| IL-1α | Naive | KD | 1.000000 | ns |
| IL-1α | Con | KD | 1.000000 | ns |
| IL-2 | Naive | Con | 0.190000 | ns |
| IL-2 | Naive | KD | 0.422000 | ns |
| IL-2 | Con | KD | 1.000000 | ns |
| IL-4 | Naive | Con | 1.000000 | ns |
| IL-4 | Naive | KD | 0.557000 | ns |
| IL-4 | Con | KD | 0.559000 | ns |
| IL-6 | Naive | Con | 0.000197 | *** |
| IL-6 | Naive | KD | 0.011800 | * |
| IL-6 | Con | KD | 0.000468 | *** |
| IL-7 | Naive | Con | 0.003070 | ** |
| IL-7 | Naive | KD | 0.024700 | * |
| IL-7 | Con | KD | 0.000943 | *** |
| IL-9 | Naive | Con | 1.000000 | ns |
| IL-9 | Naive | KD | 0.283000 | ns |
| IL-9 | Con | KD | 0.124000 | ns |
| KC | Naive | Con | 0.000478 | *** |
| KC | Naive | KD | 0.004200 | ** |
| KC | Con | KD | 0.003370 | ** |
| LIX | Naive | Con | 0.009750 | ** |
| LIX | Naive | KD | 1.000000 | ns |
| LIX | Con | KD | 0.010700 | * |
| Leptin | Naive | Con | 1.000000 | ns |
| Leptin | Naive | KD | 1.000000 | ns |
| Leptin | Con | KD | 0.940000 | ns |
| Lymphotactin | Naive | Con | 1.000000 | ns |
| Lymphotactin | Naive | KD | 0.073200 | ns |
| Lymphotactin | Con | KD | 0.045800 | * |
| MCP-1 | Naive | Con | 0.198000 | ns |
| MCP-1 | Naive | KD | 0.149000 | ns |
| MCP-1 | Con | KD | 1.000000 | ns |
| MCSF | Naive | Con | 1.000000 | ns |
| MCSF | Naive | KD | 0.538000 | ns |
| MCSF | Con | KD | 1.000000 | ns |
| MIG | Naive | Con | 0.165000 | ns |
| MIG | Naive | KD | 0.053500 | ns |
| MIG | Con | KD | 0.013200 | * |
| MIP-1α | Naive | Con | 0.043700 | * |
| MIP-1α | Naive | KD | 0.050800 | ns |
| MIP-1α | Con | KD | 1.000000 | ns |
| MIP-1γ | Naive | Con | 0.000692 | *** |
| MIP-1γ | Naive | KD | 0.001040 | ** |
| MIP-1γ | Con | KD | 0.222000 | ns |
| RANTES | Naive | Con | 0.001950 | ** |
| RANTES | Naive | KD | 0.005080 | ** |
| RANTES | Con | KD | 0.077100 | ns |
| SDF-1 | Naive | Con | 0.008490 | ** |
| SDF-1 | Naive | KD | 0.020400 | * |
| SDF-1 | Con | KD | 0.293000 | ns |
| TCA-3 | Naive | Con | 0.753000 | ns |
| TCA-3 | Naive | KD | 1.000000 | ns |
| TCA-3 | Con | KD | 0.425000 | ns |
| TECK | Naive | Con | 1.000000 | ns |
| TECK | Naive | KD | 1.000000 | ns |
| TECK | Con | KD | 1.000000 | ns |
| TIMP-1 | Naive | Con | 0.148000 | ns |
| TIMP-1 | Naive | KD | 0.977000 | ns |
| TIMP-1 | Con | KD | 0.408000 | ns |
| TIMP-2 | Naive | Con | 1.000000 | ns |
| TIMP-2 | Naive | KD | 0.060600 | ns |
| TIMP-2 | Con | KD | 0.065300 | ns |
| TNF-α | Naive | Con | 0.398000 | ns |
| TNF-α | Naive | KD | 1.000000 | ns |
| TNF-α | Con | KD | 1.000000 | ns |
| sTNF RI | Naive | Con | 0.008120 | ** |
| sTNF RI | Naive | KD | 0.013700 | * |
| sTNF RI | Con | KD | 0.675000 | ns |
| sTNF RII | Naive | Con | 0.003750 | ** |
| sTNF RII | Naive | KD | 0.003050 | ** |
| sTNF RII | Con | KD | 1.000000 | ns |
